# High Genetic Diversity and Adaptive Potential of Two Simian Hemorrhagic Fever Viruses in a Wild Primate Population

**DOI:** 10.1101/001040

**Authors:** Adam L. Bailey, Michael Lauck, Andrea Weiler, Samuel D. Sibley, Jorge M. Dinis, Zachary Bergman, Chase W. Nelson, Michael Correll, Michael Gleicher, David Hyeroba, Alex Tumukunde, Geoffrey Weny, Colin Chapman, Jens H. Kuhn, Austin L. Hughes, Thomas C. Friedrich, Tony L. Goldberg, David H. O’Connor

## Abstract

Key biological properties such as high genetic diversity and high evolutionary rate enhance the potential of certain RNA viruses to adapt and emerge. Identifying viruses with these properties in their natural hosts could dramatically improve disease forecasting and surveillance. Recently, we discovered two novel members of the viral family *Arteriviridae*: simian hemorrhagic fever virus (SHFV)-krc1 and SHFV-krc2, infecting a single wild red colobus (*Procolobus rufomitratus tephrosceles*) in Kibale National Park, Uganda. Nearly nothing is known about the biological properties of SHFVs in nature, although the SHFV type strain, SHFV-LVR, has caused devastating outbreaks of viral hemorrhagic fever in captive macaques. Here we detected SHFV-krc1 and SHFV-krc2 in 40% and 47% of 60 wild red colobus tested, respectively. We found viral loads in excess of 10^6^ − 10^7^ RNA copies per milliliter of blood plasma for each of these viruses. SHFV-krc1 and SHFV-krc2 also showed high genetic diversity at both the inter- and intra-host levels. Analyses of synonymous and non-synonymous nucleotide diversity across viral genomes revealed patterns suggestive of positive selection in SHFV open reading frames (ORF) 5 (SHFV-krc2 only) and 7 (SHFV-krc1 and SHFV-krc2). Thus, these viruses share several important properties with some of the most rapidly evolving, emergent RNA viruses.

## INTRODUCTION

Certain RNA viruses have biological properties that make them particularly likely to emerge [1]. High genetic diversity, high evolutionary rates, and high viral loads are all thought to enhance the potential of some RNA viruses to adapt to changing environments by evading immune responses within hosts or enabling the invasion of new host populations [2,3]. It is widely accepted that identifying and characterizing such viruses in their natural hosts is important for disease monitoring and prevention [4–7]. For example, the origin of human immunodeficiency virus (HIV)-1, group M (the strain responsible for the AIDS pandemic) from simian immunodeficiency viruses (SIVs) of wild chimpanzees in Central Africa [8] underscores the importance of “pandemic prevention,” as well as the importance of non-human primates as reservoirs of potentially important viruses.

The simian hemorrhagic fever viruses (SHFVs) are a poorly understood group of single stranded, positive-sense RNA viruses within the family *Arteriviridae* that have only recently been detected in wild primates [9,10]. Almost everything known about these viruses comes from the type strain of simian hemorrhagic fever virus (SHFV-LVR), which caused several “explosive” disease outbreaks in captive macaques (*Macaca assamensis, M. arctoides, M. fasciularis, M. nemestrina, and M. mulatta*) between 1964 and 1996. [11–14]. The lethality of SHFV infection in these Asian Old World monkeys (OWMs) suggested that macaques were highly susceptible to the virus, and were therefore unlikely to be natural hosts of SHFV-LVR. Further investigation revealed that monkeys of several African OWM species – specifically patas monkeys (*Erythrocebus patas*), grivets (*Chlorocebus aethiops*), and Guinea baboons (*Papio papio*) – could persistently harbor SHFV-LVR in captivity without signs of disease [15]. Although this finding implicated African OWMs as the immediate source of SHFV-LVR in the captive outbreaks, neither SHFV-LVR nor any of its relatives had ever been identified in a wild animal until recently [9,16].

In 2011, we discovered two highly divergent simian arteriviruses infecting a single wild red colobus (*Procolobus rufomitratus tephrosceles*) in Kibale National Park, Uganda (hereafter Kibale), which we named SHFV-krc1 and SHFV-krc2 [9]. Subsequently, we discovered additional, highly divergent simian arteriviruses in red-tailed guenons (*Cercopithecus ascanius*) from the same location [10]. Here we characterize SHFV-krc1 and SHFV-krc2 in 60 red colobus from Kibale. We show that these viruses infect a high proportion of red colobus in this population, replicate to high titers in infected monkeys, and have high genetic diversity, both within and among hosts. Our findings demonstrate that these viruses possess properties that are associated with the rapid evolutionary adaptability characteristic of many emerging RNA viruses.

## MATERIALS AND METHODS

### Arterivirus genome organization and nomenclature

SHFV genomes contain a duplication of four open reading frames (ORFs) relative to the other viruses in the *Arteriviridae* family: porcine reproductive and respiratory syndrome virus (PRRSV), equine arteritis virus (EAV), and lactate dehydrogenase-elevating virus of mice (LDV). Previous publications regarding SHFV have treated the naming of these additional ORFs inconsistently. For clarity, we have adopted the nomenclature scheme presented in [17], and have included a schematic (Figure 1) to maintain continuity with previous publications.

**Figure 1.**
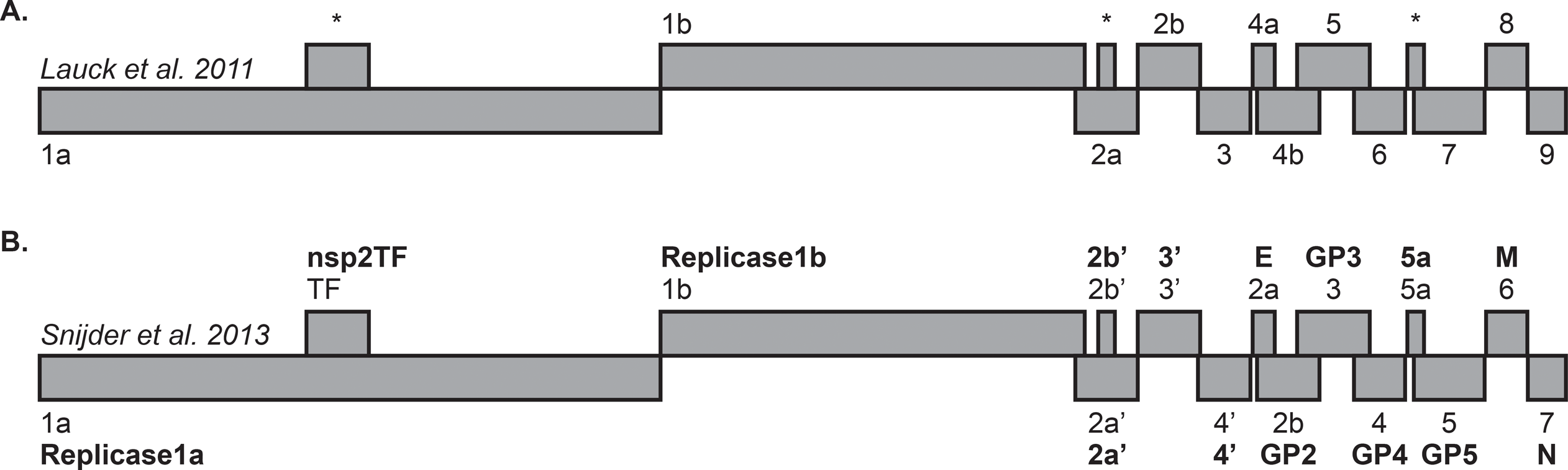
Schematic of the SHFV genome. (A) ORFs as they are referred to in Lauck et al., 2011 [9], labeled sequentially 5’-3’: ORF1a-ORF9. Asterisks denote ORFs identified in SHFV-krc1 and SHFV-krc2 not reported in Lauck et al., 2011 [9]. (B) ORFs as they are named in Snijder et al., 2013 [17], labeled 5’-3’: ORF1a-ORF7, with duplicated ORFs designated by a “prime” (*e.g.* ORF2a’). Expression products are given in bold.

### Ethics statement

All animal use in this study followed the guidelines of the Weatherall Report on the use of non-human primates in research. Specific protocols were adopted to minimize suffering through anesthesia and other means during capture, immobilization, and sampling of the non-human primates. These included use of anesthesia during capture (Ketamine/Xylazine, administered intramuscularly with a variable-pressure pneumatic rifle), minimization of immobilization time and the use of an anesthetic reversal agent (Atipamezole) to reduce recovery time, and conservative limits on blood sample volumes (<1% body weight), as previously described [9]. Following sampling, all animals were released back to their social group without incident [18]. All research was approved by the Uganda Wildlife Authority (permit UWA/TDO/33/02), the Uganda National Council for Science and Technology (permit HS 364), and the University of Wisconsin Animal Care and Use Committee (protocol V01409-0-02-09) prior to initiation of the study.

### Study site and sample collection

Red colobus were sampled between 2/5/2010 and 7/22/2012 in Kibale National Park, Uganda, a 795 km^2^ semi-deciduous park in western Uganda (0°13’-0°41’ N, 30°19’-30°32’ E) known for its exceptional density of primates belonging to diverse species. Blood was separated using centrifugation and plasma was frozen immediately in liquid nitrogen for storage and transport to the United States. Samples were shipped in an IATA-approved dry shipper to the USA for further analysis at the Wisconsin National Primate Research Center in accordance with CITES permit #002290 (Uganda).

### Molecular methods

Samples were processed for sequencing in a biosafety level 3 laboratory as described previously [9,10]. Briefly, for each animal, one ml of blood plasma was filtered (0.45 µm) and viral RNA was isolated using the Qiagen QIAamp MinElute virus spin kit (Qiagen, Hilden, Germany), omitting carrier RNA. DNase treatment was performed and cDNA synthesis was accomplished using random hexamers. Samples were fragmented and sequencing adaptors were added using the Nextera DNA Sample Preparation Kit (Illumina, San Diego, CA, USA). Deep sequencing was performed on the Illumina MiSeq (Illumina, San Diego, CA, USA).

### SHFV detection by quantitative RT-PCR

We developed a multiplex quantitative RT-PCR (qRT-PCR) assay to quantify plasma viral RNA of both SHFV-krc1 and SHFV-krc2 in each sample. Taqman assays were designed with amplification primers specific for either SHFV-krc1 (5’- ACACGGCTACCCTTACTCC-3’ and 5’- TCGAGGTTAARCGGTTGAGA-3’) or SHFV-krc2 (5’-AACGCGCACCAACCACTATG-3’ and 5’GCGTGTTGAGGCCCTAATTTG-3’). The SHFV-krc1 probe (5’-Quasar 670- TTCTGGTCCTCTTGCGAAGGC-BHQ2-3’) and SHFV-krc2 probe (5’-6-Fam-TTTGCTCAAGCCAATGACCTGCG-BHQ1-3’) were also virus-specific. The fluorophores used do not produce overlapping spectra, so no color compensation was required. Viral RNA was reverse transcribed and quantified using the SuperScript III One-Step qRT-PCR system (Invitrogen, Carlsbad, CA) on a LightCycler 480 (Roche, Indianapolis, IN). Reverse transcription was carried out at 37°C for 15 min and then 50°C for 30 min followed by two minutes at 95°C and 50 cycles of amplification as follows: 95°C for 15 sec and 60°C for 1 min. The reaction mixture contained MgSO^4^ at a final concentration of 3.0 mM, 150 ng random primers (Promega, Madison, WI), with all 4 amplification primers at a concentration of 600 nM and both probes at a concentration of 100 nM.

### Genetic analyses

Sequence data were analyzed using CLC Genomics Workbench 5.5 (CLC bio, Aarhus, Denmark) and Geneious R5 (Biomatters, Auckland, New Zealand). Low quality (<Q25) and short reads (<100 bp) were removed and the full genome sequences for each virus were acquired using *de novo* assembly. Due to the approximately 52% nucleotide sequence similarity between the genomes of SHFVkrc1 and SHFV-krc2, and the high frequency of co-infections in our animal cohort, we devised a method to minimize mapping of SHFV-krc1 reads to SHFV-krc2 (and vice versa) within a co-infected animal. Briefly, total reads from a co-infected animal were mapped to the SHFV-krc1 consensus sequence generated from *de novo* assembly and “unmapped reads” were collected, then mapped to the SHFV-krc2 consensus sequence obtained from *de novo* assembly. The resulting SHFV-krc2 consensus sequence was then used as the reference for mapping and collecting unmapped reads to map to the SHFV-krc1 consensus sequence generated from *de novo* assembly. This process was repeated until changes between the reference and the consensus sequences were not observed for either virus. Using this method, reads corresponding to SHFV-krc1 and SHFV-krc2 were reliably segregated in co-infected animals, with less than 0.2% of SHFV-specific reads mapping to both viruses. The average coverage per genome was 5,654x (range 118-19,115x) for SHFV-krc1 variants and 2,264 (range 94-6,613x) for SHFV-krc2 variants. For intra-host genetic analysis, sequencing reads were mapped to the corresponding consensus sequence for each variant. Single nucleotide polymorphism (SNP) reports were generated in Geneious, with a minimum coverage threshold of 100 reads and a minimum frequency threshold of five percent.

### Evolutionary analyses

The synonymous nucleotide diversity (π_S_) and the non-synonymous nucleotide diversity (π_N_) were estimated for each ORF individually from SNP reports generated by mapping sequencing reads to their corresponding consensus sequence. We estimated π_S_ = n_s_/L_s_ and π_N_ = n_n_/L_n_, where n_s_ is the mean number of pairwise synonymous differences; n_n_ is the mean number of pairwise synonymous differences; L_s_ is the number of synonymous sites; and L_n_ is the number of nonsynymous sites. L_s_ and L_n_ were estimated by the method described in [19]. To compare viruses across different hosts, variant consensus sequences were aligned by the CLUSTAL algorithm in MEGA 5.05 [20]. Estimating π_S_ and π_N_ separately for each ORF in each virus from co-infected animals, we used a factorial analysis of variance to test for main effects of the virus (SHFV-krc1 vs. SHFV-krc2) and the ORF, and for virus-by-ORF interactions. In the case of π_S_, there were highly significant main effects of virus (F_1, 459_ = 41.31; *p* < 0.001) and of ORF (F_13, 459_ = 14.07; *p* < 0.001), but there was not a significant virus-by-ORF interaction (F_13, 459_ = 1.35; n.s.). In the case of π_N_, there were significant main effects of virus (F_1, 459_ = 4.42; *p* = 0.036) and of ORF (F_13, 459_ = 53.26; *p* < 0.001), and there was a highly significant virus-by-ORF interaction (F_13, 459_ = 4.39; *p* < 0.001). Sliding window analysis involved estimating π_S_ and π_N_ in a sliding window of 9 codons, numbered according to the numbering in the sequence alignment of the first codon in the window.

### Layercake visualization

We developed a specialized visualization tool called LayerCake for this dataset. This tool allows visual comparison of variants for multiple individuals simultaneously, encoding sequences as bands of color, with redder sections of the band corresponding to regions with a higher proportion of polymorphic reads. Downloadable versions of the krc1 and krc2 datasets are available, along with a generalized tutorial for interpreting LayerCake displays, at http://graphics.cs.wisc.edu/Vis/LayerCake/.

## RESULTS

### Sample collection and infection frequency of SHFV-krc1 and SHFV-krc2 in Kibale red colobus

Blood samples were collected from 60 adult red colobus residing in the Kanyawara area of Kibale over a period of 2.5 years. These animals represent approximately half of a defined social group, but comprise a relatively small proportion of the total red colobus population in Kibale [21]. All animals appeared normal and healthy at the time of sampling. RNA was isolated from the blood plasma of each animal and “unbiased” deep sequencing was performed on an Illumina MiSeq machine as previously described [9,10]. *De novo* assembly and iterative mapping of sequencing reads yielded 52 near full-length SHFV consensus sequences (GenBank accession numbers KC787607-KC787658). Twenty-four animals (40.0%) were infected with SHFV-krc1, and 28 animals (46.7%) were infected with SHFV-krc2. Twenty-one animals (35.0%) were co-infected with both SHFV-krc1 and SHFV-krc2 (Figure 2).

**Figure 2.**
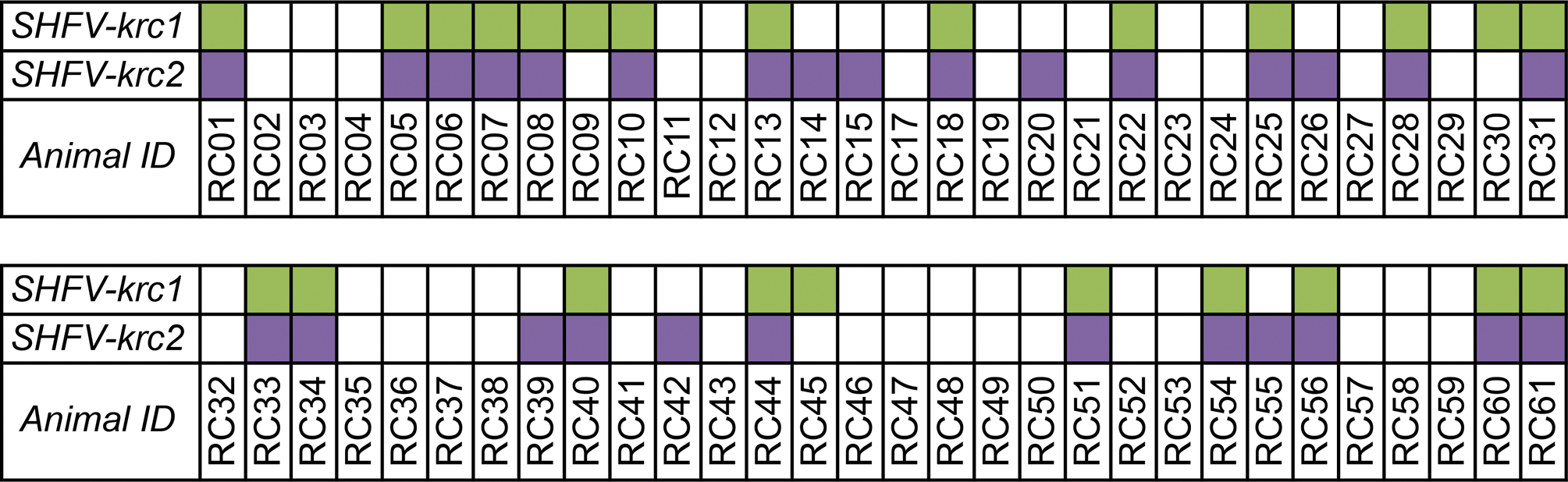
Infection of the Kibale red colobus with SHFV-krc1 and SHFV-krc2. SHFV-krc1 (green) and SHFV-krc2 (purple) infections were identified by “unbiased” deep sequencing and confirmed by strain-specific qRT-PCR.

### Viral loads of SHFV-krc1 and SHFV-krc2 in the Kibale red colobus

To estimate the viral load of SHFV-krc1 and SHFV-krc2 in infected red colobus, a strain-specific qRT-PCR assay was designed to amplify highly conserved regions in ORF7 of the SHFV-krc1 and SHFV-krc2 genomes. This assay was used to assess the viral burden in cell-free plasma for each animal found to be positive by deep sequencing. SHFV-krc1 viremia was consistently high, averaging 5.1 × 10^7^ vRNA copies/ml plasma, (range: 1.5 × 10^6^ − 1.9 × 10^8^ copies/ml plasma) (Figure 3A). SHFV-krc2 loads were more varied (range: 3.4 × 10^4^ − 4.1 × 10^7^ copies/ml) and significantly lower than SHFV-krc1 with an average plasma titer of 7.5 × 10^6^ vRNA copies/ml plasma (*p* = 0.0001, two-tailed t-test). Although instances of mono-infection were scarce relative to co-infection, mono/co-infection status did not impact the load of either virus to a statistically significant extent (mono- vs. co-infected: *p* = 0.063 for SHFV-krc1, *p* = 0.089 for SHFV-krc2, two-tailed t-test, Figure 3B,C).

**Figure 3.**
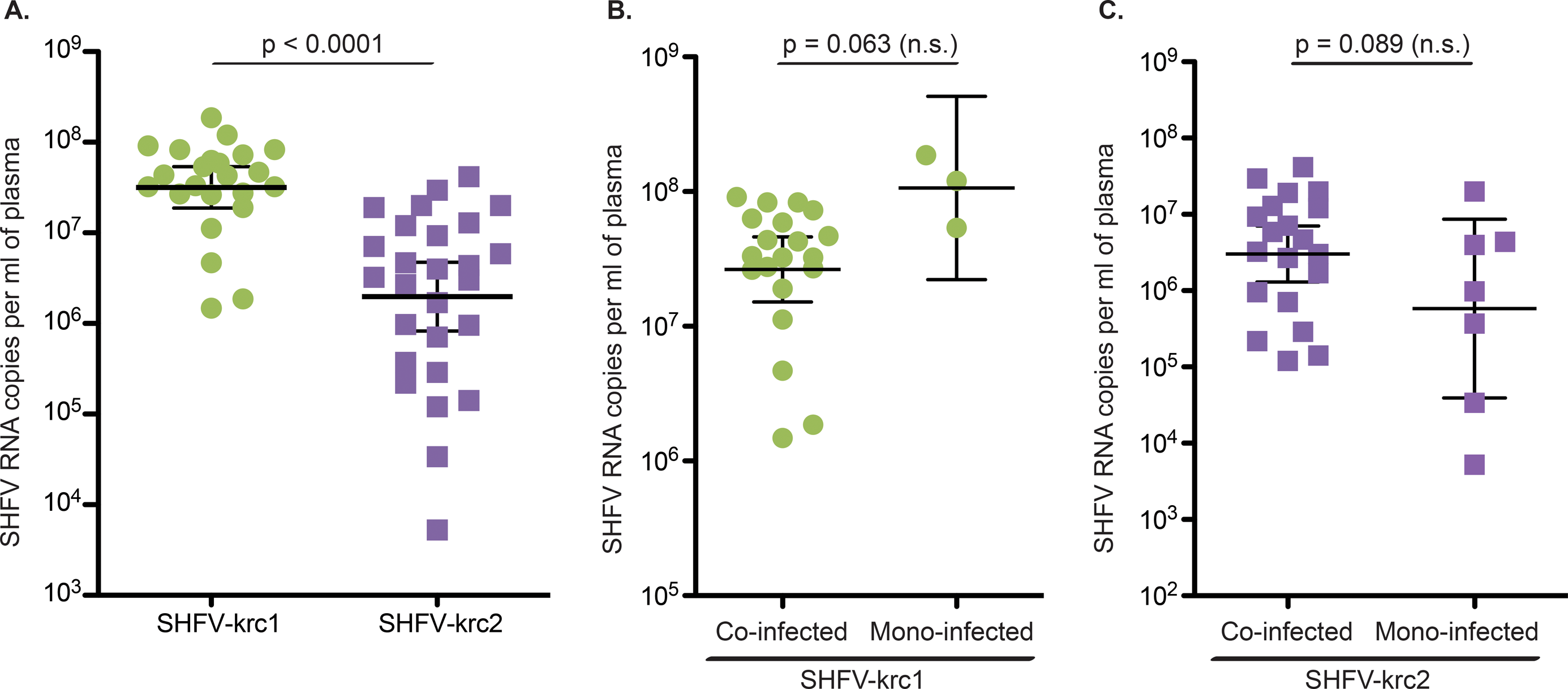
Viral loads of SHFV-krc1 and SHFV-krc2 in the Kibale red colobus. Comparison of SHFV-krc1 (green) and SHFV-krc2 (purple) viral loads from all animals positive for either virus (A) and viral loads from mono-infections vs. co-infections of SHFV-krc1 (B) and SHFV-krc2 (C). RNA was isolated from blood plasma and quantitative RT-PCR was performed using strain-specific primers and probes designed from deep sequencing data. Statistical significance was assessed using a two-tailed t-test performed on log-transformed values (CI = 95%).

### Consensus-level genetic diversity among SHFV-krc1 and SHFV-krc2 variants

To quantify the genetic diversity of SHFV-krc1 and SHFV-krc2 within the red colobus population, we examined similarity among the 24 SHFV-krc1 and 28 SHFV-krc2 variants by comparing the nucleotide consensus sequences of each viral variant (Figure 4). These consensus sequences represent the majority nucleotide base present at each position of the genome for the viral population within each host. Because RNA viruses often exist within a host as a highly heterogeneous population (*i.e. “*mutant swarm”), the consensus sequence may not actually be present in the within-host viral population [22, 23]. Nevertheless, the construction of consensus sequences allowed us to compare the average viral population of each variant (*i.e*. inter-host diversity). For SHFV-krc1, percent pairwise nucleotide identity between variants ranged from 86.9%–99.5%. A highly related core group (SHFV-krc1 from red colobus 06, 28, 33, 22, 25, 34, 54, 31, 40, 05, 56, 44, 08, 45, 01) with pairwise nucleotide identities ranging from 94.5%–99.3% comprised 63% of the variants (Figure 4A). A distinct second group (SHFV-krc1 from red colobus 09, 30, 10, 18, 07, 60) with a slightly wider range of similarity (92.0%–99.5% pairwise nucleotide identity) made up an additional 21% of variants. A similar pattern, with two distinct groups, was observed for SHFV-krc2 variants (range: 89.67%–99.48% pairwise nucleotide identity, Figure 4B). However, patterns of SHFV-krc1 genetic similarity among hosts were different from patterns of SHFV-krc2 similarity among hosts. (data not shown). Interestingly, SHFV-krc1 variants from red colobus 13 and 61 were highly dissimilar to all other SHFV-krc1 variants identified, with pairwise nucleotide identities ranging from 86.8%–88.7%.

**Figure 4.**
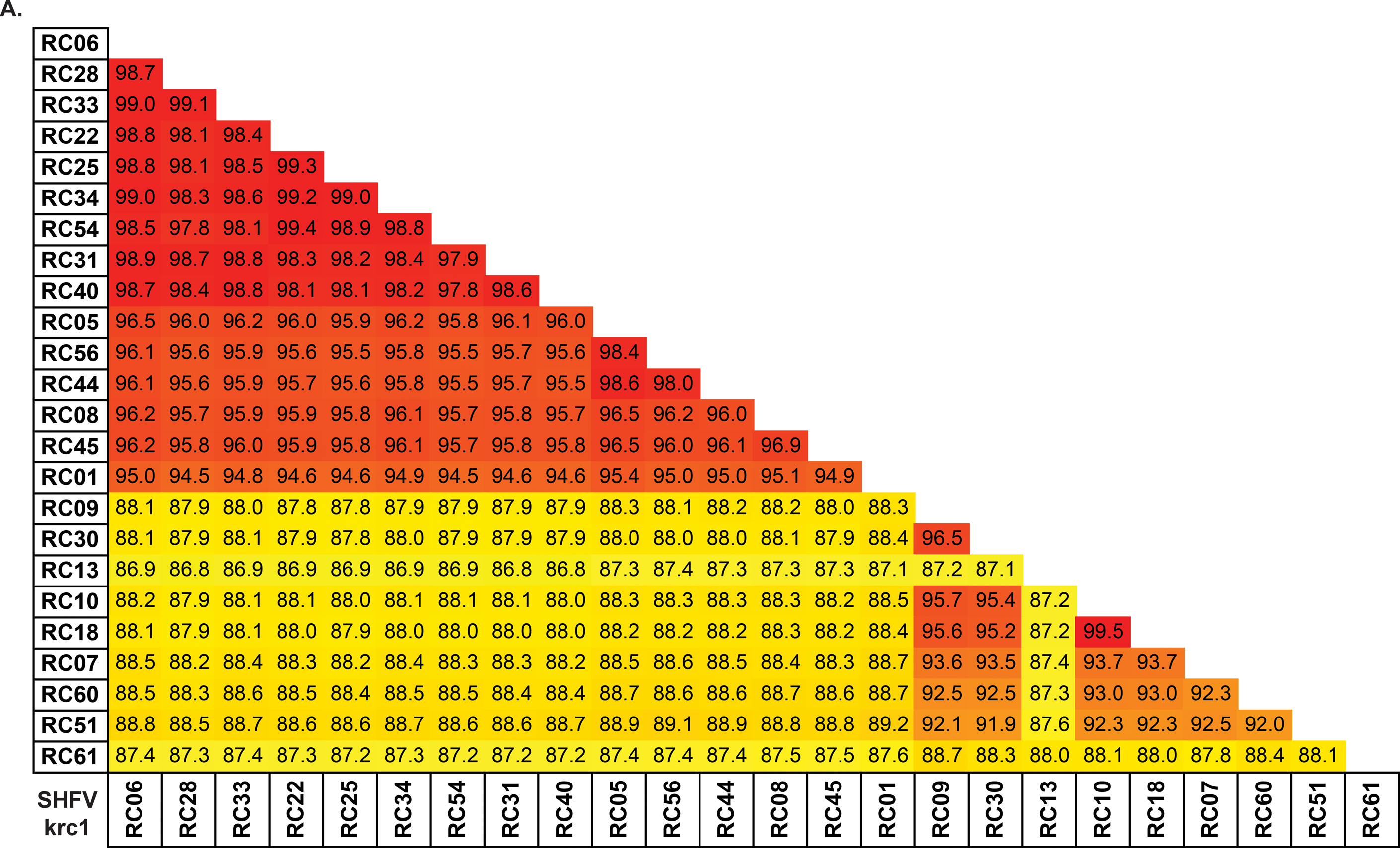
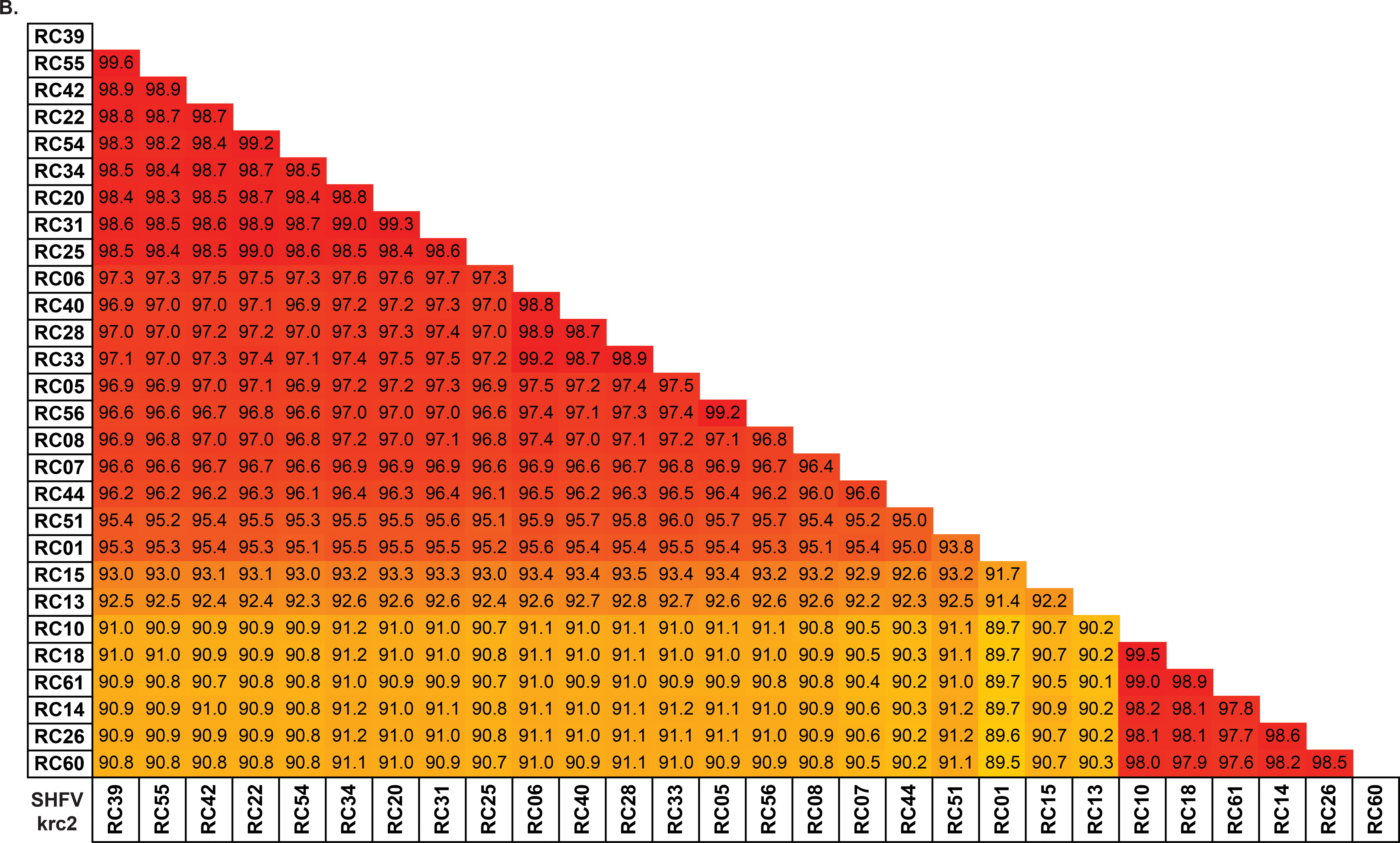
Pairwise comparison of nucleotide identity among variants of SHFV-krc1 and SHFV-krc2 from Kibale red colobus (RC). Full coding sequences for each isolate were aligned using CLC Genomics Workbench. Numbers show percent nucleotide identity between two variants within (A) SHFV-krc1 or (B) SHFV-krc2. Colors highlight similarity, with red representing the most similar sequences and yellow representing sequences with the lowest degree of nucleotide identity. The same color scale was used for (A) and (B).

### Within-host genetic diversity of SHFV-krc1 and SHFV-krc2

To examine the genetic diversity of SHFV-krc1 and SHFV-krc2 within individual monkeys, we calculated the non-synonymous and synonymous nucleotide diversity, π_N_ and π_S_ respectively, for each within-host viral population using deep sequencing reads from each viral variant. Comparing π_N_ and π_S_ from specific regions of a viral genome can reveal the mode of natural selection acting on a region. For example, π_N_ < π_S_ is indicative of negative selection acting to remove deleterious protein-coding mutations, while π_N_ > π_S_ is suggestive of positive selection acting to drive beneficial protein-coding mutations to fixation. We found that, overall, negative selection acting against deleterious non-synonymous mutations predominated for both SHFV-krc1 and SHFV-krc2. In SHFV-krc1, π_S_ exceeded π_N_ by a ratio of over 6:1, whereas in SHFV-krc2, π_S_ exceeded π_N_ by a ratio of nearly 5:1. Both π_S_ and π_N_ were significantly greater in SHFV-krc1 than in SHFV-krc2 (*p* = 0.002 and *p* = 0.021, paired t-test), indicating greater overall nucleotide diversity in SHFV-krc1 than in SHFV-krc2 (Figure 5). A positive correlation between viral load and both π_S_ and π_N_ was observed. However, mean π_S_ and π_N_ did not differ significantly between co-infected monkeys and those infected with only SHFV-krc1 or SHFV-krc2 (data not shown).

**Figure 5.**
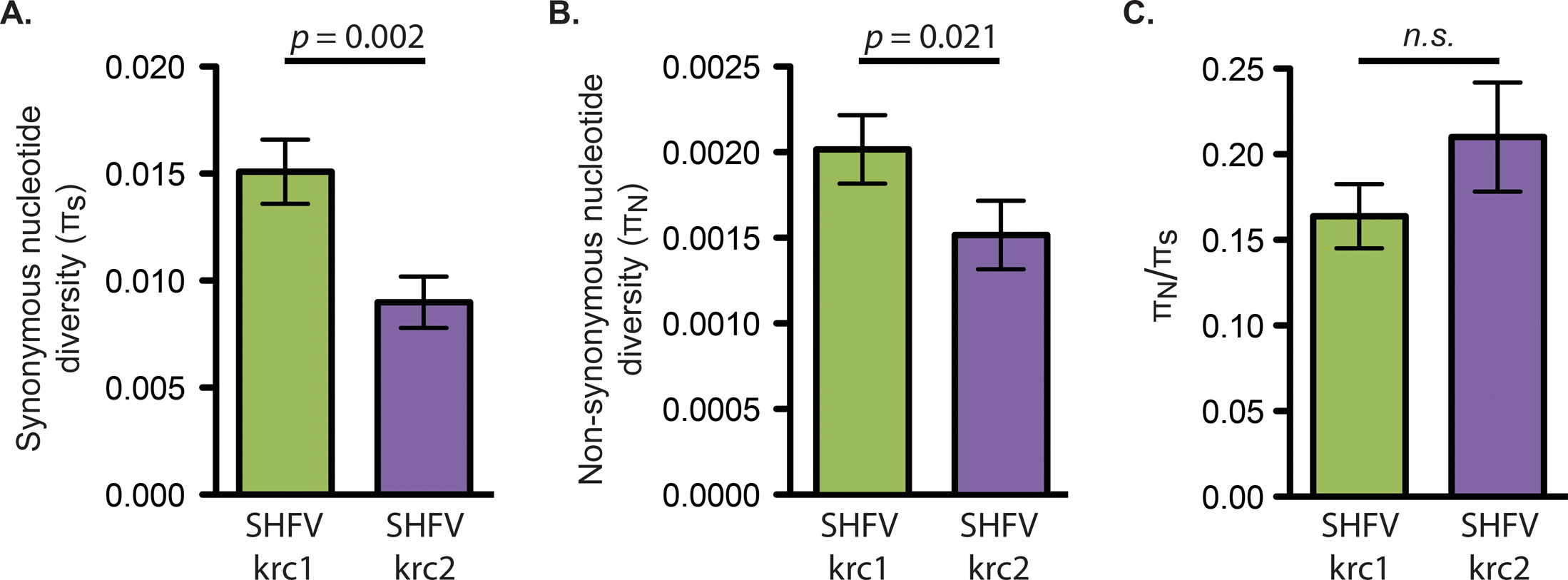
Overall nucleotide diversity of SHFV-krc1 and SHFV-krc2. Mean (± S.E.) π_S_ (A), π_N_ (B), and π_N_/π_S_ (C) in monkeys infected with SHFV-krc1 (green) and SHFV-krc2 (purple). Paired t-tests were performed to compare mean values between SHFV-krc1 and SHFV-krc2.

The organization of ORFs in the genomes of SHFV-krc1 and SHFV-krc2 was the same as described previously (Figure 1) [9,10,17], so we used a factorial analysis of variance approach to investigate π_S_ and π_N_ in ORFs in both viruses. In general, 3’-proximal ORFs displayed more non-synonymous diversity than 5’-proximal ORFs, suggesting that the proteins encoded by 5’-proximal ORFs may be more functionally constrained than those encoded by 3’-proximal ORFs. However, the extent to which underlying RNA structures may have affected this analysis is unknown [24–26]. ORF5 showed the highest mean π_N_ in SHFV-krc1 and among the highest in SHFV-krc2 (Figure 6). In the case of both SHFV-krc1 and SHFV-krc2, a sliding window plot of 9 codons revealed peaks of π_N_ corresponding to codons 1-46 and 64-100 of ORF5 (Figure 7A,B). The latter peak (codons 64-100) also involved high π_S_, suggesting a mutational hotspot. Interestingly, π_N_ was substantially higher in ORF3 of SHFV-krc2 than of SHFV-krc1 (Figure 6). Sliding window analysis revealed a substantial peak of π_N_ between codons 141-173 of SHFV-krc2 ORF3 (Figure 7C) that greatly exceeded π_S_, suggesting strong positive selection in this region of SHFV-krc2. This peak of π_N_ corresponded to a region of variable length rich in acidic residues. An analogous peak of π_N_ in ORF3 of SHFV-krc1 was not found, although a unique peak of π_N_ was identified between codons 50 and 68 (Figure 7D). Of note, a high degree of variability in predicted *N*-glycosylation [27] was associated with each instance of elevated π_N_ in ORF3 and ORF5 for both SHFV-krc1 and SHFV-krc2. For peaks of π_N_ found in regions of ORF3 and ORF5 that shared sequence with an overlapping alternative ORF, sliding window plot analysis in the alternative ORFs revealed peaks of π_S_ demonstrating that observed elevations in π_N_ were ORF-specific, as expected [28,29] (data not shown).

**Figure 6.**
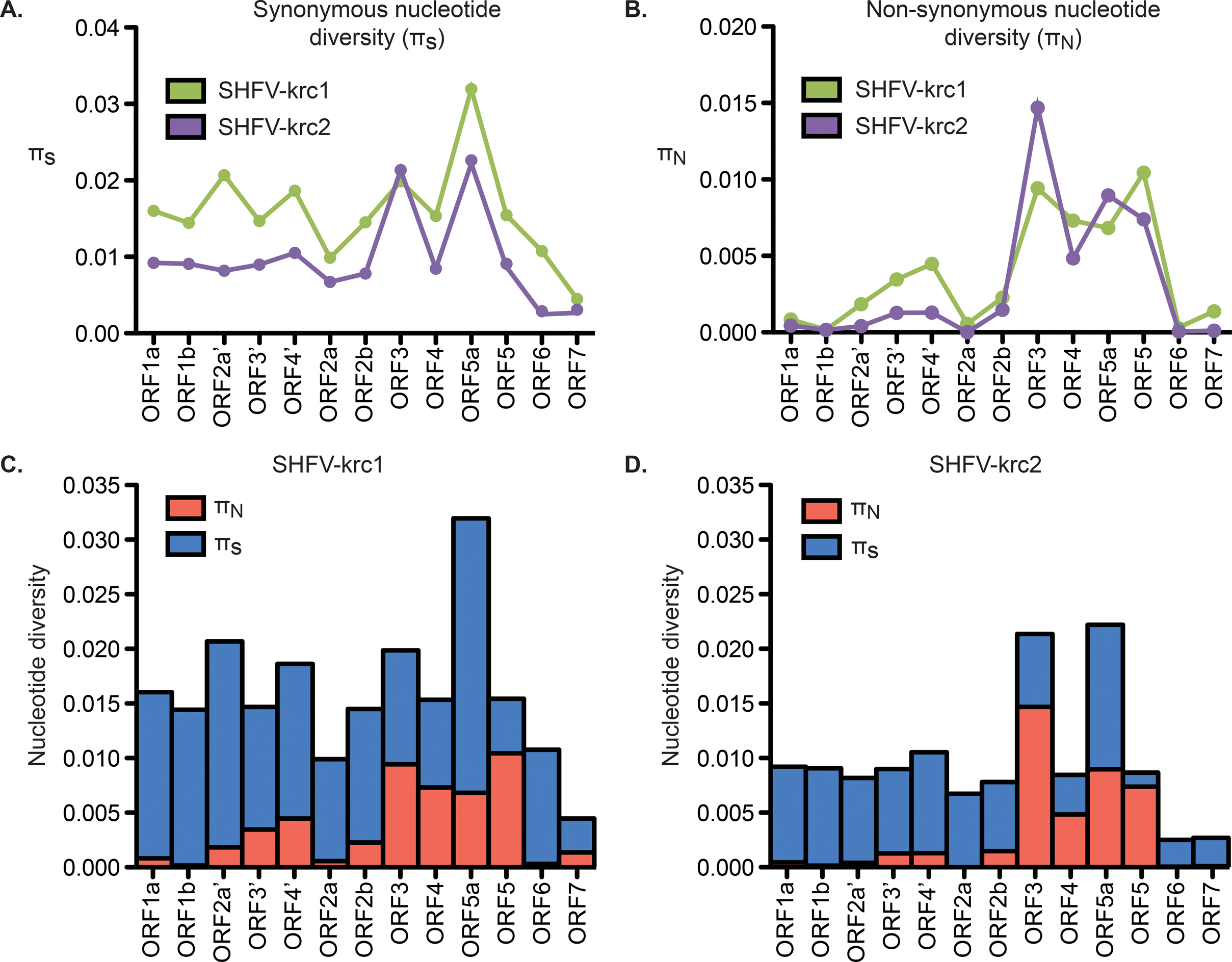
Nucleotide diversity SHFV-krc1 and SHFV-krc2 ORFs. Interaction graphs comparing mean π_S_ (A) and π_N_ (B) in ORFs from SHFV-krc1 (green) and SHFV-krc2 (purple). In the case of π_N_ there was a significant ORF-by-virus interaction (F13, 459 = 4.39; p < 0.001). Comparison of mean π_s_ (blue) to π_N_ (red) within ORFs of SHFV-krc1 (C) and SHFV-krc2 (D) revealed substantial differences among ORFs within each virus.

**Figure 7.**
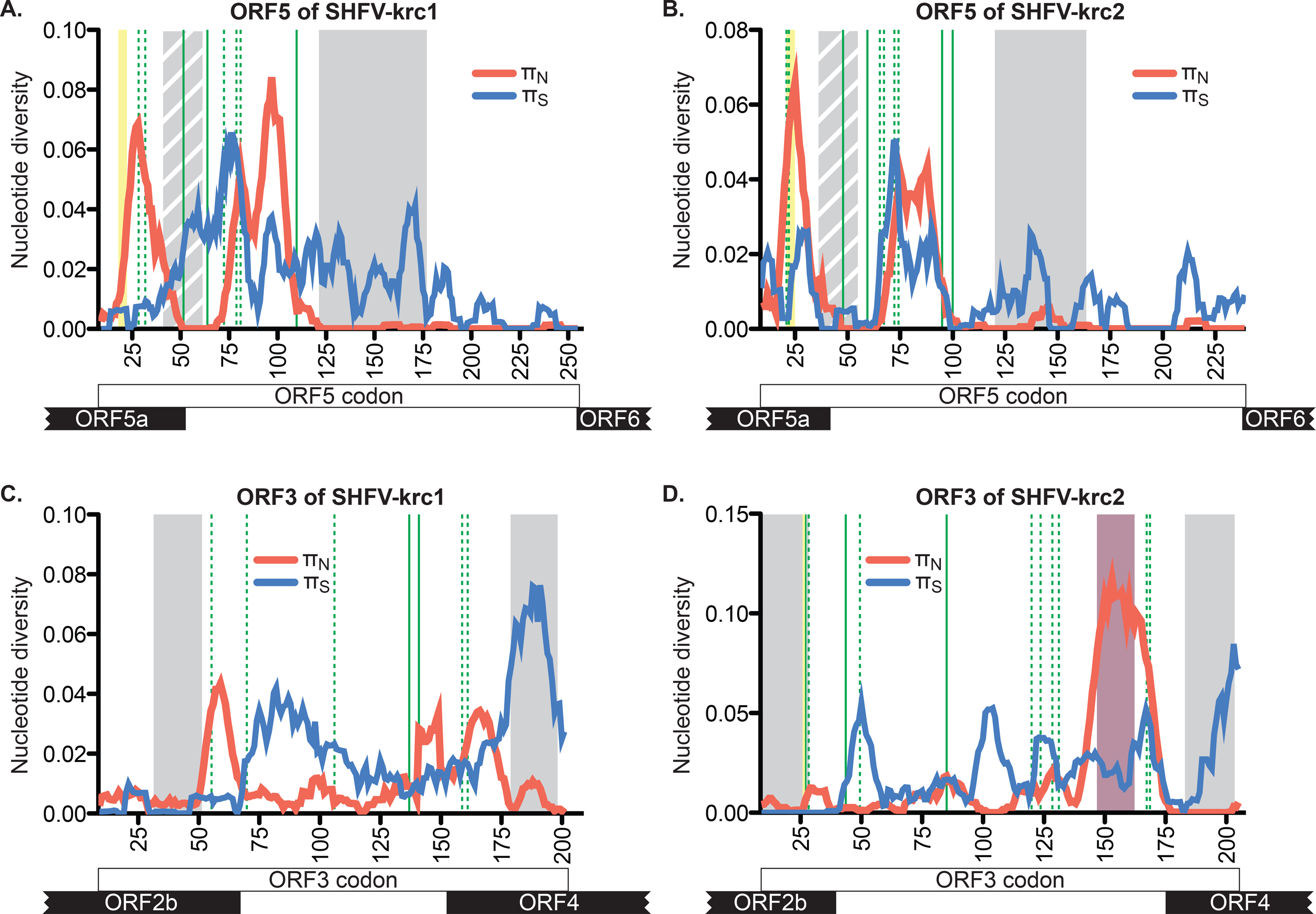
Nucleotide diversity across ORF5 and ORF7 of SHFV-krc1 and SHFV-krc2. Mean π_S_ (blue) and π_N_ (red) in sliding windows of 9 codons across the coding region of ORF5 (A,B) and ORF3 (C,D). Overlapping ORFs are shown at the bottom. Grey boxes represent predicted transmembrane domains, with striped grey boxes representing a hydrophobic region unique to the SHFVs. Green lines depict putative sites of *N*-glycosylation, with dashed green lines showing sites that are variably glycosylated. Yellow boxes show predicted signal peptide cleavage sites that vary in location in GP5 of SHFV-krc1 and SHFV-krc2 and were not found in GP3 of SHFV-krc1. The purple box corresponds to the unique region of highly variable acidic residues found only in ORF3 of SHFV-krc2.

Unique patterns of inter- and intra-host variation can be visualized on a genome-wide scale for all SHFV-krc1 and SHFV-krc2 variants using our custom-built LayerCake software: http://graphics.cs.wisc.edu/Vis/LayerCake/.

## DISCUSSION

This study provides the first systematic analysis of SHFV genetic diversity in a population of wild non-human primates. Our findings show that SHFV-krc1 and SHFV-krc2 have a high frequency of infection in the red colobus population of Kibale, and that these viruses achieve high titers in the blood of infected monkeys. Our study also details, for the first time, the genetic diversity of SHFV-krc1 and SHFV-krc2 both within and among infected hosts. We draw particular attention to the signatures of natural selection identified throughout the genomes of these viruses, with an emphasis on signatures of positive selection identified in ORFs 3 and 5.

To date, primates from only two species – the red colobus and red-tailed guenon – have been found to harbor simian arteriviruses in the wild [9, 10]. However, the origins and host-ranges of these viruses are not clear. Our findings support the hypothesis that simian arteriviruses are endemic to African OWMs and cause little to no clinical disease in these hosts. However, when introduced into Asian OWMs, these viruses may be lethal, as exemplified by SHFV-LVR [13,30]. This pattern of pathogenesis is similar to SIV [31] and, like SIV, the simian arteriviruses appear to be well host-adapted, which suggests an ancient evolutionary relationship between these viruses and their African OWM hosts. This is in contrast to the arterivirus PRRSV, which emerged suddenly in pig populations across the globe in the 1980’s [32]. Taken together, this implies that the prevalence and diversity of the *Arteriviridae*, including the simian arterivirus group, may be greater than currently appreciated.

SHFV-krc1 and SHFV-krc2 display many biological properties associated with the potential for rapid evolution – a feature that is shared by many emergent RNA viruses. For example, high diversity at the population level (inter-host diversity) can facilitate speciation, and related yet distinct viruses can recombine [31,33]. High within-host diversity also enables a virus to escape the host immune response, alter tropism, and infect new host species [34,35]. In these contexts, high viral load increases the probability of transmission by “widening” the population bottleneck that often reduces the fitness of an RNA virus upon transmission [36–38]. Such features enhance the ability of a virus to adapt to changing environments and have been implicated in the ability of some viruses to transmit across species barriers [2]. Although the arteriviruses in general are considered to be highly specific for their hosts, we note that SHFV-LVR and related viruses have been transmitted between primate species from presumptive African primate hosts into Asian macaques on several occasions [11–14, 30]. Recent work suggests that the capacity for SHFVs to infect multiple primate species is not unique to SHFV-LVR, as experimental infection of macaques with SHFV-krc1 resulted in viral replication and clinical disease (unpublished data). The biological properties of SHFV-krc1 and SHFV-krc2 in a natural host that we have identified herein may help explain the propensity of the SHFVs to infect primates of species other than their natural host. Future investigation of these viruses should provide further insight into the full extent of their cross-species transmission potential.

Our analysis shows that SHFV-krc1 and SHFV-krc2 are not merely highly divergent forms of the same virus, but in fact possess unique and distinct biological properties. Nucleotide diversity was consistently higher in SHFV-krc1 than in SHFV-krc2. This is likely a result of the higher viral loads observed for SHFV-krc1, reflecting more extensive viral replication and a correspondingly higher rate of accumulation of within-host mutations [39]. This hypothesis is supported by positive correlations between viral load and both synonymous and non-synonymous nucleotide diversity (data not shown). Interestingly, viral load and nucleotide diversity for both SHFV-krc1 and SHFV-krc2 were not significantly impacted by the presence of the other virus (Figure 3). When viewed in light of the competitive exclusion principle [40] this finding suggests that the two viruses may occupy discrete niches within the red colobus host (*e.g.* tissue tropisms), possibly resulting in distinct aspects of infection that could contribute to the observed differences in infection frequency (Figure 2) and viral burden (Figure 3).

The most significant difference in nucleotide diversity that we observed between SHFV-krc1 and SHFV-krc2 was found in ORF3 (Figures 6 and 7), which codes for the putative envelope glycoprotein GP3. GP3 of SHFV-krc1 and SHFV-krc2 appears similar in topology to GP3 of other arteriviruses, with predicted N- and C-terminal membrane-spanning domains separated by a heavily glycosylated ectodomain. While the precise function of GP3 in the arterivirus life-cycle remains elusive, GP3 is thought to be an important determinant of tissue tropism [41,42]. GP3 is also immunogenic [43,44] and glycans attached to the GP3 ectodomain may play a role in evasion of the humoral immune response through the shielding neutralizing antibody epitopes [45]. It is possible that GP3 has multiple functions, as GP3 of PRRSV and LDV have been found in both virion-associated and soluble secreted forms [46–50]. Our analysis revealed a distinct region of non-synonymous diversity suggestive of positive selection in ORF3 of SHFV-krc2 (codons 141-173) (Figure 7D). This region contained an unusually high density of acidic residues and multiple, variable putative *N*-glycosylation sites. Although a similar region was not found in ORF3 of SHFV-krc1, a unique peak of non-synonymous diversity was identified between codons 50–68 of ORF3 in SHFV-krc1 that was also suggestive of positive selection. Finally, another difference between SHFV-krc1 and SHFV-krc2 was that no signal sequence cleavage site could be identified in GP3 of any SHFV-krc1 variant, while a clear signal sequence cleavage site was found C-terminal to the first predicted transmembrane domain in GP3 of SHFV-krc2 [51]. The most likely explanation of this finding is that the signal sequence cleavage site of GP3 in SHFV-krc2 is not utilized, as has been shown for GP3 of EAV [43,49]. Although the functional significance of these differences between GP3 of SHFV-krc1 and GP3 of SHFV-krc2 is presently unclear, the potential effect of GP3 on the immune response and its putative role as a determinant of host cell tropism may help to explain why SHFV-krc2 mono-infections are twice as common as SHFV-krc1 mono-infections (Figure 2), despite the significantly lower viral loads of SHFV-krc2 relative to SHFV-krc1 (Figure 3A).

Despite the differences we observed between SHFV-krc1 and SHFV-krc2 in ORF3, we found nearly identical patterns of non-synonymous and synonymous nucleotide diversity in ORF5, which – by analogy to other arteriviruses - codes for the major envelope glycoprotein GP5 [17,52]. Two distinct peaks of non-synonymous diversity were found in the 5’-proximal region of ORF5, which corresponds to the protein’s predicted ectodomain (Figure 7). This region of GP5 contains the primary neutralizing antibody epitope of PRRSV, EAV, and LDV [53–56], as well as an immunodominant “decoy” epitope in PRRSV that may serve to subvert neutralizing antibody responses [57]. These epitopes align closely with more 3’-proximal peak of non-synonymous diversity we identified in SHFV-krc1 and SHFV-krc2 (data not shown), suggesting that antibody pressure in the red colobus may select for escape mutations in SHFV-krc1 and SHFV-krc2, resulting in the observed genetic diversity of this region.

Glycans in this region of the GP5 ectodomain – in addition to aiding viral attachment through the binding of host molecules (*e.g.* sialoadhesin for PRRSV) [58] - are also implicated in evasion of humoral immune responses by arteriviruses. Pigs infected with PRRSV variants containing partially de-glycosylated GP5 mount significantly more robust neutralizing antibody responses [45,59]. A similar observation was made for LDV in mice, and the abolishment of *N*-glycosylation sites in GP5 had the additional effect of altering the tissue tropism of these “neurotropic” LDV strains [60,61]. Putative *N*-glycosylation sites were variably found in association with each peak of non-synonymous nucleotide diversity identified ORF5/GP5 of both SHFV-krc1 and SHFV-krc2 (Figure 7). However, in contrast to the GP5 ectodomains of PRRSV, EAV, and LDV, a highly conserved hydrophobic stretch of approximately thirty amino acids separated these two regions of diversity, and was predicted to form an additional transmembrane domain in both SHFV-krc1 and SHFV-krc2 [62–64]. A domain that spans the membrane once in this region would place the N-terminal portion of GP5 – including the region corresponding to the more 5’-proximal peak of non-synonymous nucleotide diversity – within the virion. While this possibility cannot be formally excluded, the high sequence diversity of this region – including multiple putative *N*-glycosylation sites – suggests that this scenario is unlikely. Nevertheless, it is conceivable that this region interacts extensively with the membrane of the virion and its functional significance, although obscure, is highlighted by its conservation across all other known simian arteriviruses including SHFV-LVR, SHFV-krtg1, and SHFV-krtg2 (data not shown).

The findings presented in this study show that SHFV variants contain high genetic diversity within their hosts. This presents the possibility that SHFV-krc1 or SHFV-krc2 could evolve rapidly within the red colobus, perhaps gaining virulence, similar to the recent emergence of highly pathogenic PRRSV in pigs in China and Southeast Asia [65,66]. As the red colobus population of Kibale faces the stressors of deforestation and a changing climate, monitoring these infections may be important to the conservation of this already endangered wild primate [67].

## REFERENCES

1. Holmes EC, Grenfell BT (2009) Discovering the phylodynamics of RNA viruses. PLoS computational biology 5: e1000505.

2. Parrish CR, Holmes EC, Morens DM, Park E-C, Burke DS et al (2008) Cross-species virus transmission and the emergence of new epidemic diseases. Microbiology and molecular biology reviews : MMBR 72: 457–470.

3. Grenfell BT, Pybus OG, Gog JR, Wood JLN, Daly JM et al (2004) Unifying the epidemiological and evolutionary dynamics of pathogens. Science (New York, NY) 303: 327–332.

4. King DA, Peckham C, Waage JK, Brownlie J, Woolhouse MEJ (2006) Epidemiology. Infectious diseases: preparing for the future. Science (New York, NY) 313: 1392–1393.

5. Woolhouse M, Scott F, Hudson Z, Howey R, Chase-Topping M (2012) Human viruses: discovery and emergence. Philosophical transactions of the Royal Society of London Series B, Biological sciences 367: 2864–2871.

6. Morse SS, Mazet JAK, Woolhouse M, Parrish CR, Carroll D et al (2012) Prediction and prevention of the next pandemic zoonosis. Lancet 380: 1956–1965.

7. Wolfe ND, Dunavan CP, Diamond J (2007) Origins of major human infectious diseases. Nature 447: 279–283.

8. Sharp PM, Hahn BH (2011) Origins of HIV and the AIDS Pandemic. Cold Spring Harbor perspectives in medicine 1: a006841.

9. Lauck M, Hyeroba D, Tumukunde A, Weny G, Lank SM et al (2011) Novel, divergent simian hemorrhagic fever viruses in a wild Ugandan red colobus monkey discovered using direct pyrosequencing. PloS one 6: e19056.

10. Lauck M, Sibley SD, Hyeroba D, Tumukunde A, Weny G et al (2013) Exceptional simian hemorrhagic Fever virus diversity in a wild african primate community. Journal of virology 87: 688–691.

11. Lapin BA, Pekerman SM, Iakovleva LA, Dzhikidze EK, Shevtsova ZV et al (1967) [Hemorrhagic fever in monkeys]. Voprosy virusologii 12: 168–173.

12. Allen AM, Palmer AE, Tauraso NM, Shelokov A (1968) Simian hemorrhagic fever. II. Studies in pathology. The American journal of tropical medicine and hygiene 17: 413–421.

13. Palmer AE, Allen AM, Tauraso NM, Shelokov A (1968) Simian hemorrhagic fever. I. Clinical and epizootiologic aspects of an outbreak among quarantined monkeys. The American journal of tropical medicine and hygiene 17: 404–412.

14. Tauraso NM, Shelokov A, Palmer AE, Allen AM (1968) Simian hemorrhagic fever. 3. Isolation and characterization of a viral agent. The American journal of tropical medicine and hygiene 17: 422–431.

15. London WT (1977) Epizootiology, transmission and approach to prevention of fatal simian haemorrhagic fever in rhesus monkeys. Nature 268: 344–345.

16. Gravell M, London WT, Leon ME, Palmer AE, Hamilton RS (1986) Differences among isolates of simian hemorrhagic fever (SHF) virus. Proceedings of the Society for Experimental Biology and Medicine Society for Experimental Biology and Medicine (New York, NY) 181: 112–119.

17. Snijder EJ, Kikkert M, Fang Y (2013) Arterivirus molecular biology and pathogenesis. The Journal of general virology 94: 2141–2163.

18. Wasserman MD, Chapman CA, Milton K, Goldberg TL, Ziegler TE (2013) Physiological and Behavioral Effects of Capture Darting on Red Colobus Monkeys (Procolobus rufomitratus) with a Comparison to Chimpanzee (Pan troglodytes) Predation. International Journal of Primatology 34: 1020–1031.

19. Nei M, Gojobori T (1986) Simple methods for estimating the numbers of synonymous and nonsynonymous nucleotide substitutions. Molecular biology and evolution 3: 418–426.

20. Tamura K, Peterson D, Peterson N, Stecher G, Nei M et al (2011) MEGA5: molecular evolutionary genetics analysis using maximum likelihood, evolutionary distance, and maximum parsimony methods. Molecular biology and evolution 28: 2731–2739.

21. Goldberg TL, Paige S, Chapman CA (2012) The Kibale EcoHealth Project: exploring connections among human health, animal health, and landscape dynamics in western Uganda. In: Aguirre AA, Daszak P, Ostfeld RS, editors. Conservation Medicine: Applied Cases of Ecosystem Health. New York: Oxford University Press.

22. Holmes EC (2009) The Evolution and Emergence of RNA Viruses. Oxford: Oxford University Press.

23. Wright CF, Morelli MJ, Thebaud G, Knowles NJ, Herzyk P et al (2011) Beyond the consensus: dissecting within-host viral population diversity of foot-and-mouth disease virus by using next-generation genome sequencing. Journal of virology 85: 2266–2275.

24. Brierley I (1995) Ribosomal frameshifting viral RNAs. The Journal of general virology 76: 1885–1892.

25. van Marle G, Dobbe JC, Gultyaev AP, Luytjes W, Spaan WJ et al (1999) Arterivirus discontinuous mRNA transcription is guided by base pairing between sense and antisense transcription-regulating sequences. Proceedings of the National Academy of Sciences of the United States of America 96: 12056–12061.

26. Fang Y, Treffers EE, Li Y, Tas A, Sun Z et al (2012) Efficient-2 frameshifting by mammalian ribosomes to synthesize an additional arterivirus protein. Proceedings of the National Academy of Sciences of the United States of America 109: E2920–8.

27. Zhang M, Gaschen B, Blay W, Foley B, Haigwood N et al (2004) Tracking global patterns of N-linked glycosylation site variation in highly variable viral glycoproteins: HIV, SIV, and HCV envelopes and influenza hemagglutinin. Glycobiology 14: 1229–1246.

28. de Oliveira T, Salemi M, Gordon M, Vandamme A-M, van Rensburg EJ et al (2004) Mapping sites of positive selection and amino acid diversification in the HIV genome: an alternative approach to vaccine design? Genetics 167: 1047–1058.

29. Hughes AL, Westover K, da Silva J, O’Connor DH, Watkins DI (2001) Simultaneous positive and purifying selection on overlapping reading frames of the tat and vpr genes of simian immunodeficiency virus. Journal of virology 75: 7966–7972.

30. Johnson RF, Dodd LE, Yellayi S, Gu W, Cann JA et al (2011) Simian hemorrhagic fever virus infection of rhesus macaques as a model of viral hemorrhagic fever: clinical characterization and risk factors for severe disease. Virology 421: 129–140.

31. Apetrei C, Robertson DL, Marx PA (2004) The history of SIVS and AIDS: epidemiology, phylogeny and biology of isolates from naturally SIV infected non-human primates (NHP) in Africa. Frontiers in bioscience: a journal and virtual library 9: 225–254.

32. Wensvoort G, Terpstra C, Pol JM, ter Laak EA, Bloemraad M et al. (1991) Mystery swine disease in The Netherlands: the isolation of Lelystad virus. The Veterinary quarterly 13: 121–130.

33. Snijder EJ, Meulenberg JJ (1998) The molecular biology of arteriviruses. The Journal of general virology 79: 961–979.

34. Farci P, Shimoda A, Coiana A, Diaz G, Peddis G et al (2000) The outcome of acute hepatitis C predicted by the evolution of the viral quasispecies. Science (New York, NY) 288: 339–344.

35. Wolinsky SM, Korber BT, Neumann AU, Daniels M, Kunstman KJ et al (1996) Adaptive evolution of human immunodeficiency virus-type 1 during the natural course of infection. Science (New York, NY) 272: 537–542.

36. Novella IS, Duarte EA, Elena SF, Moya A, Domingo E et al (1995) Exponential increases of RNA virus fitness during large population transmissions. Proceedings of the National Academy of Sciences of the United States of America 92: 5841–5844.

37. Keele BF, Giorgi EE, Salazar-Gonzalez JF, Decker JM, Pham KT et al (2008) Identification and characterization of transmitted and early founder virus envelopes in primary HIV-1 infection. Proceedings of the National Academy of Sciences of the United States of America 105: 7552–7557.

38. Bordería AV, Lorenzo-Redondo R, Pernas M, Casado C, Alvaro T et al (2010) Initial fitness recovery of HIV-1 is associated with quasispecies heterogeneity and can occur without modifications in the consensus sequence. PloS one 5: e10319.

39. Kimura M (1983) The Neutral Theory of Molecular Evolution. Cambridge: Cambridge University Press.

40. Armstrong RA, McGehee R (1980) Competitive Exclusion. The American Naturalist 115: 151–170.

41. Tian D, Wei Z, Zevenhoven-Dobbe JC, Liu R, Tong G et al (2012) Arterivirus minor envelope proteins are a major determinant of viral tropism in cell culture. Journal of virology 86: 3701–3712.

42. Lu Z, Zhang J, Huang CM, Go YY, Faaberg KS et al (2012) Chimeric viruses containing the N-terminal ectodomains of GP5 and M proteins of porcine reproductive and respiratory syndrome virus do not change the cellular tropism of equine arteritis virus. Virology 432: 99–109.

43. Hedges JF, Balasuriya UB, MacLachlan NJ (1999) The open reading frame 3 of equine arteritis virus encodes an immunogenic glycosylated, integral membrane protein. Virology 264: 92–98.

44. Plana-Duran J, Climent I, Sarraseca J, Urniza A, Cortes E et al (1997) Baculovirus expression of proteins of porcine reproductive and respiratory syndrome virus strain Olot/91. Involvement of ORF3 and ORF5 proteins in protection. Virus genes 14: 19–29.

45. Vu HLX, Kwon B, Yoon K-J, Laegreid WW, Pattnaik AK et al (2011) Immune evasion of porcine reproductive and respiratory syndrome virus through glycan shielding involves both glycoprotein 5 as well as glycoprotein 3. Journal of virology 85: 5555–5564.

46. Faaberg KS, Plagemann PG (1997) ORF 3 of lactate dehydrogenase-elevating virus encodes a soluble, nonstructural, highly glycosylated, and antigenic protein. Virology 227: 245–251.

47. Mardassi H, Gonin P, Gagnon CA, Massie B, Dea S (1998) A subset of porcine reproductive and respiratory syndrome virus GP3 glycoprotein is released into the culture medium of cells as a non-virion-associated and membrane-free (soluble) form. Journal of virology 72: 6298–6306.

48. Gonin P, Mardassi H, Gagnon CA, Massie B, Dea S (1998) A nonstructural and antigenic glycoprotein is encoded by ORF3 of the IAF-Klop strain of porcine reproductive and respiratory syndrome virus. Archives of virology 143: 1927–1940.

49. Wieringa R, de Vries AAF, Raamsman MJB, Rottier PJM (2002) Characterization of two new structural glycoproteins, GP(3) and GP(4), of equine arteritis virus. Journal of virology 76: 10829–10840.

50. de Lima M, Ansari IH, Das PB, Ku BJ, Martinez-Lobo FJ et al (2009) GP3 is a structural component of the PRRSV type II (US) virion. Virology 390: 31–36.

51. Petersen TN, Brunak S, von Heijne G, Nielsen H (2011) SignalP 4.0: discriminating signal peptides from transmembrane regions. Nature methods 8: 785–786.

52. Dokland T (2010) The structural biology of PRRSV. Virus research 154: 86–97.

53. Balasuriya UB, Patton JF, Rossitto PV, Timoney PJ, McCollum WH et al (1997) Neutralization determinants of laboratory strains and field isolates of equine arteritis virus: identification of four neutralization sites in the amino-terminal ectodomain of the G(L) envelope glycoprotein. Virology 232: 114–128.

54. Plagemann PG (2001) Complexity of the single linear neutralization epitope of the mouse arterivirus lactate dehydrogenase-elevating virus. Virology 290: 11–20.

55. Plagemann PGW, Rowland RRR, Faaberg KS (2002) The primary neutralization epitope of porcine respiratory and reproductive syndrome virus strain VR-2332 is located in the middle of the GP5 ectodomain. Archives of virology 147: 2327–2347.

56. Plagemann PGW (2004) The primary GP5 neutralization epitope of North American isolates of porcine reproductive and respiratory syndrome virus. Veterinary immunology and immunopathology 102: 263–275.

57. Ostrowski M, Galeota JA, Jar AM, Platt KB, Osorio FA et al (2002) Identification of neutralizing and nonneutralizing epitopes in the porcine reproductive and respiratory syndrome virus GP5 ectodomain. Journal of virology 76: 4241–4250.

58. Van Breedam W, Van Gorp H, Zhang JQ, Crocker PR, Delputte PL et al (2010) The M/GP(5) glycoprotein complex of porcine reproductive and respiratory syndrome virus binds the sialoadhesin receptor in a sialic acid-dependent manner. PLoS pathogens 6: e1000730.

59. Ansari IH, Kwon B, Osorio FA, Pattnaik AK (2006) Influence of N-linked glycosylation of porcine reproductive and respiratory syndrome virus GP5 on virus infectivity, antigenicity, and ability to induce neutralizing antibodies. Journal of virology 80: 3994–4004.

60. Chen Z, Li K, Rowland RR, Plagemann PG (1999) Selective antibody neutralization prevents neuropathogenic lactate dehydrogenase-elevating virus from causing paralytic disease in immunocompetent mice. Journal of neurovirology 5: 200–208.

61. Chen Z, Li K, Plagemann PG (2000) Neuropathogenicity and sensitivity to antibody neutralization of lactate dehydrogenase-elevating virus are determined by polylactosaminoglycan chains on the primary envelope glycoprotein. Virology 266: 88–98.

62. Krogh A, Larsson B, von Heijne G, Sonnhammer EL (2001) Predicting transmembrane protein topology with a hidden Markov model: application to complete genomes. Journal of molecular biology 305: 567–580.

63. von Heijne G (1992) Membrane protein structure prediction. Hydrophobicity analysis and the positive-inside rule. Journal of molecular biology 225: 487–494.

64. K H, W S (1993) TMBASE - A database of membrane spanning protein segments. Biol Chem Hoppe-Seyler 374,166:

65. Ni J, Yang S, Bounlom D, Yu X, Zhou Z et al (2012) Emergence and pathogenicity of highly pathogenic Porcine reproductive and respiratory syndrome virus in Vientiane, Lao People’s Democratic Republic. Journal of veterinary diagnostic investigation : official publication of the American Association of Veterinary Laboratory Diagnosticians, Inc 24: 349–354.

66. Tian K, Yu X, Zhao T, Feng Y, Cao Z, (2007) Emergence of fatal PRRSV variants: unparalleled outbreaks of atypical PRRS in China and molecular dissection of the unique hallmark. PloS one 2: e526.

67. Struhsaker T (2008) Procolobus rufomitratus ssp. tephrosceles. In: IUCN 2013. IUCN Red List of Threatened Species.

